# *Apolipoprotein E* reduces the number and activation of non-invariant Natural Killer T cells

**DOI:** 10.1101/2025.07.04.663139

**Authors:** Rituparna Chakrabarti, Sushmitha Duddu, Praphulla Chandra Shukla

**Author notes:** Department of Anesthesiology and Critical Care Medicine, The Johns Hopkins University School of Medicine, Baltimore, Maryland, United States of America (Rituparna Chakrabarti). Department of Human Medicine, Carl von Ossietzky University, Oldenburg, Lower Saxony, Germany (Praphulla Chandra Shukla).

## Abstract

Natural Killer T (NKT) cells, which modulate atherosclerosis, include two groups – invariant (iNKT) and variant (vNKT). These subsets differentially regulate the disease progression. Yet, the role of vNKTs in atherosclerosis remains unclear. We induced atherosclerosis by feeding high-fat diet (HFD) to *Apoe^-/-^* and analyzed the vNKTs in the liver and spleen. The vNKTs were termed non-iNKTs since they were negatively selected within the NKT population. Available literature suggests NKTs as lipid-recognizing cells; however, to our surprise, the non-iNKT numbers and phenotype remained unchanged between HFD-fed and chow-fed *Apoe^-/-^*. This was a blindsiding and unexpected outcome of the non-iNKTs being unaltered and unaffected with or without HFD, indicating no observable impact of atherosclerosis on these subsets. Albeit remaining unperturbed by atherosclerosis, these non-iNKTs demonstrated an identical but unique increase and upregulated activation in both the chow and HFD-fed *Apoe^-/-^*. These results instigated an investigation of the baseline correlation of the non-iNKTs between young C57BL/6 (WT) and *Apoe^-/-^*. Previously unknown and confounding results revealed upregulated activation and increased non-iNKT numbers but decreased IL-4^+^ non-iNKTs in the *Apoe^-/-^* compared to WT. Furthermore, HFD-fed WT that developed dyslipidemia, elucidated increased hepatic non-iNKTs and splenic IFN-γ^+^ non-iNKTs compared to chow-fed WT controls. These results were not perceived in chow and HFD-fed *Apoe^-/-^*. Although lipid-responsive, non-iNKTs in *Apoe^-/-^* mice failed to respond to lipid stress, unlike those in dyslipidemic WT mice. These findings reveal that loss of *Apoe*, rather than atherosclerosis itself, drives altered non-iNKT biology. Thus, *Apoe* deficiency intrinsically dysregulates non-iNKTs, masking disease-associated immune changes. *Apoe* loss alters non-iNKT number and function, independent of atherosclerosis, and challenges the interpretation of immune responses by NKT subsets in *Apoe^-/-^* models. Therefore, this study warrants the use of *Apoe* null mice in studying NKT cells.

## Introduction

Atherosclerosis is a disease of the vascular walls that develops gradually with chronic dyslipidemia, activating innate and adaptive immunity. NKTs are a specific group of T cell subsets sharing the features of NK and T cells, also designated as ‘unconventional’ T cell subsets. These cells home in the liver but are also found in the spleen, blood, lymph nodes, intestine, lungs, mucosa, etc. ^1^. These cells are located in human and mice atherosclerotic plaques ^2^. NKT cells are restricted to CD1d molecules that present endogenous glycolipids to the NKTs. Thus, it becomes pertinent to perceive the participation of NKT cells in modulating the initiation and progression of atherosclerosis or discern the effect of atherosclerosis on the quantity and quality of these cells. The NKTs are subdivided into two subsets based on their T cell receptor (TCR) composition: the invariant (iNKT) or the type I and the variant (vNKT) or the type IIs. As the name suggests, the invariant or the iNKTs have an invariant TCR, and the variant or the vNKTs have diverse TCRs ^3, 4^. Previous reports designate NKTs and the invariant subset as pro-inflammatory cells participating in atherosclerosis ^5–7^. The absence of the whole NKT cells and only the iNKT subset deficiency ameliorated atherosclerosis ^8^. However, reports from other groups, emphasize how the NKT cells suppress atherosclerosis ^9, 10^. Furthermore, contrasting results from Kritikou *et al.* suggest that mast cells with *CD1d* deficiency aggravate atherosclerosis also supporting the anti-atherosclerotic nature of NKT cells ^11^. In our meta-analysis, we analyzed various NKT cell-based studies in atherosclerosis and reported that both the whole NKT cells and the iNKT subset promote atherosclerosis. Most of these studies are performed on atheroprone compound knockout animals that are deficient in either NKT or only the iNKT cells on the background of either *Apoe* or *Ldlr* gene (*CD1d^-/-^; Apoe^-/-^* or *CD1d^-/-^; Ldlr^-/-^*, *Jα18^-/-^; Apoe^-/-^* or *Jα18^-/-^; Ldlr^-/-^*) ^6, 9, 11–20^. These studies were conducted using atheroprone mice because despite feeding HFD to WT mice, they do not develop atherosclerosis. There are reports on the differential functioning of the T cells, macrophages, and iNKT cells in *Apoe^-/-^* mice ^21–25^. Here, this study reveals the role of the *Apoe* gene in the vNKT cells. To our knowledge, there is no data about the number and function of the vNKTs cells in atherosclerosis. This study aimed to identify the vNKT functions in atherosclerosis induced in *Apoe^-/-^*. Eventually, this study unveils the importance of the *Apoe* gene on the number and function of the vNKT subset.

## Material and methods

### Animals

All animal procedures were approved by the Institutional Animal Ethics Committee (IAEC) of the Indian Institute of Technology Kharagpur (IE-5/PS-SMST/1.19). Animals were maintained under a 12/12hr light/dark cycle, room temperature at 22-25°C, and diet and water were given ad libitum. C57BL/6 and *Apoe^-/-^* mice were weaned at 21 days of age and fed a chow diet until eight weeks of age. At 8-10 weeks, the male mice were switched to a HFD containing 1.25% added cholesterol (Research Diets) for 20 weeks. Age-matched male mice were fed with a chow diet as the control group. All experiments were carried out blind to the genotype/treatment group.

### Quantifying the Development of Atherosclerosis

At the end of 20 weeks of HFD, both chow and HFD-fed animals were sacrificed, and the basal portion of the heart and proximal aortic roots were excised and embedded in the cryo-embedding media, OCT (Leica) for cryosectioning (7μm). Hearts were sectioned horizontally towards the aortic arch. Once the tri-valve leaflets of the aortic root were identified, all the sequential sections were collected on positively charged slides. For each mouse, eight serial cryosections were stained with Oil Red O (ORO) (Sigma) and counter-stained with hematoxylin. Lesion images were acquired using a Leica DM750 microscope at 4x magnification. Atherosclerotic lesions were identified by manually tracing the ORO-positive areas and quantified by NIH ImageJ analysis software as mean plaque size.

### Serum Analysis

Following the high anesthesia (4%) to the mouse with isofluorane at the end of the experiment, blood was drawn from the heart without opening the diaphragm using a 1mL syringe and kept in a non-heparinized tube. The whole blood was allowed to clot at room temperature for 30 minutes. The serum was separated from the whole blood by centrifugation at 10000rpm for 10 minutes at 4°C. Serum concentrations of total cholesterol (TC), triglyceride (TG), high-density lipoprotein cholesterol (HDL-C), and low-density lipoprotein cholesterol (LDL-C) were determined by colorimetric assay kits (Coral Clinical Systems, India).

### Lymphocyte Isolation

Single-cell suspensions of splenocytes were prepared by pressing the spleen through a 70μM strainer and lysing the red blood cells (RBCs) with ACK lysis buffer (150mM Ammonium Chloride, 10mM Potassium hydrogen carbonate, and 0.1mM Disodium ethylenediaminetetraacetate dissolved in water, pH-7.2-7.4). Hepatic mononuclear cells (HMNCs) were isolated as discussed^26^. Briefly, the liver was perfused with 5mL of ice-cold Hanks’ Balanced Salt Solution (HBSS), chopped into small pieces and digesting with 0.5% Collagenase type IV and protease inhibitor cocktail for 30 minutes at 37°C. Subsequently, it was passed through steel mesh and 70μM strainer for single-cell suspension and overlayed on 33% Normo-osmotic Percoll ^26^, followed by centrifugation at 800g for 30 minutes at room temperature with brake-off. The RBCs were lysed by ACK lysis buffer. Single cell suspensions were prepared from the aorta of the HFD fed mice after enzymatic digestion (125 U/mL Collagenase type XI, 60 U/mL Hyaluronidase type 1-s, 60 U/mL DNase I, and 450 U/mL Collagenase type I) of the aorta and passing through 70μM strainer ^27^.

### In vitro stimulation for intracellular cytokine staining

Aliquotes of 1×10^6^ cells/sample were prepared from the spleen, liver, and aortic lesions into U-bottom 96-well plates for staining. Splenocytes and HMNCs were stimulated with a mix of 50ng/mL PMA and 1μg/mL Ionomycin for 6 hours. Monensin was added for the last 4 hours.

### Flow Cytometry and Intracellular Cytokine Staining

Cells were aliquoted into 1×10^6^ cells/sample and incubated with LIVE/DEAD^TM^ fixable green dye (Invitrogen™) in PBS for 30 minutes on ice. After washing, cells were CD16/32 FcR blocked by incubating with purified 2.4G2 antibody, specific for FCγII/III receptors (BD Fc Block^TM)^ for 15 minutes at 4°C. Fluorescently labeled antibodies were used for staining the NKT cells for identification and phenotyping. The following is the fluorophore labeled antibody panel: APC-Cy7-anti-CD3e (Clone - 145-2C11, Cat. No. - 557596), PE-anti-NK1.1 (Clone - PK136, Cat. No. - 557391), PerCP-Cy5.5-anti-CD4 (Clone - RM4-5, Cat. No. - 550954), V500-anti-CD8a (Clone -2H7, Cat. No. - 560776), BV605-anti-CD69 (Clone - H1.2F3, Cat. No. - 563290), BV421-anti-IFN-γ (Clone - XMG1.2, Cat. No. - 563376), PE-Cy7-anti-IL-4 (Clone - 11B11, Cat. No. - 560699), APC-labelled-α-GC-loaded CD1d-tetramers (mCD1d/PBS-57; provided by NIH Tetramer Core Facility, Atlanta, GA, USA). Except for the tetramer, all the antibodies were procured from BD Pharmingen™. The antibody cocktail was prepared in PBS supplemented with 1% FBS and 50μL BD Horizon™ Brilliant Stain Buffer (563794). The cells were fixed and permeabilized with BD Cytofix/Cytoperm™ PlusFixation/Permeabilization Solution (555028) for intracellular cytokine staining. Stained cells were subjected to flow cytometry using the BD LSR Fortessa SORP model and the data was analyzed by FlowJo v10 software (BD Bioscience). Unstained, single-stained controls and fluorescence minus one were used to calculate compensation and set up the gates. A minimum of 200,000 events were analyzed per sample.

### Statistical Analysis

To obtain reliable experimental data, multiple biological replicates were performed. GraphPad Prism (v8) was used for all the statistical analysis. All data were presented as Mean ± SEM. Two-tailed Student’s t-test with nonparametric Mann-Whitney tests was conducted to compare the statistical difference between the means of the two groups. One-way analysis of variance (ANOVA) with Dunn’s multiple comparison post-test was performed to evaluate the difference between multiple groups,. Statistical significance was expressed as following - \**p* < 0.05, \*\**p* < 0.01, \*\*\**p* < 0.001, and \*\*\*\**p* < 0.0001.

## Results

### Identification of non-iNKT cells in the WT mice by negative selection strategy in flow cytometry

We developed a method of identifying non-iNKTs (vNKTs) by a negative selection flow cytometry strategy due to the lack of a standard method to identify these cells in mice ^28^. Although there are some older reports of identifying vNKT cells using the sulfatide-loaded CD1d tetramers, we believe that only a fraction of the vNKT TCRs recognize the sulfatide among the whole vNKT subset due to their heterogeneous population with diverse TCR ^28–31^, and hence failed to capture the total variant population. However, Zhou *et al*. recently published a strategic flow cytometry panel to identify the vNKT cells in human blood and liver. Yet, identification of these cells in mice remains to be accomplished ^32^. A heterogeneous subset of NKT cells that do not stain positive for iNKT cell markers falls into the vNKT category. We have followed using the term non-iNKT throughout the article from here. The non-iNKT cells are defined as CD3^+^NK1.1^+^α-GC/CD1d-tetramer^-^ (CD3^+^NK1.1^+^CD1d^-^) within the CD3^+^NK1.1^+^ NKT cell gate (Fig. 1). The non-iNKT cells do not bind to the α-GC loaded CD1d-tetramer. Therefore, the population negative for the tetramer within the NKT cell gate is identified as non-iNKTs. We have identified this cell population in the liver and spleen of WT mice (Fig. 1). This flow cytometry panel and gating strategy identified the non-iNKT cells along with the iNKT and NKT subset within the CD3^+^ T lymphocyte population. In the liver (n=5) of 8-10 weeks old WT mice, 36.88±3.96% are vNKTs, and 62.2±3.98% are iNKTs, and conversely, in the spleen (n=7), 67.3±3.28% are vNKTs and 31.71±3.32% are iNKTs (Fig. 1). The difference in the percentage of the non-iNKTs between the homing organ liver and peripheral lymphoid organ spleen corroborates with the previous reports that these cells actively participate in maintaining immune homeostasis in various organs differently ^33^.

**Figure 1.**
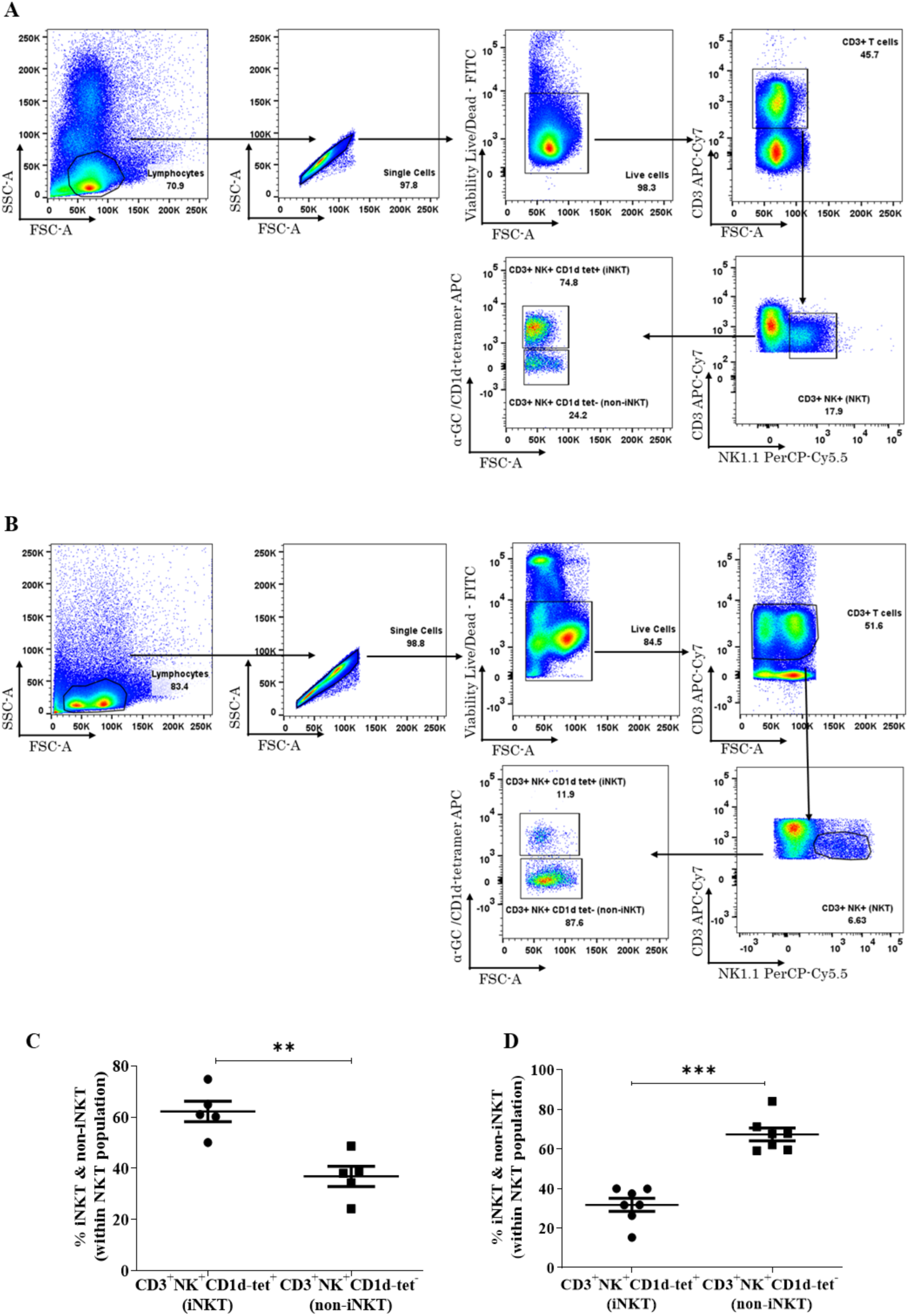
Flow cytometry gating strategy for identification of mice non-iNKT cells. Non-iNKT cells were identified within (A) HMNCs and (B) Splenocytes. The non-iNKTs were defined as CD3^+^NK1.1^+^CD1d^-^ within CD3^+^NK1.1^+^ NKT cell population. (C) Scatter plots representing the numbers of the iNKT and the non-iNKT cells in liver and spleen within the NKT cell population. All data are represented as Mean ± SEM. Statistical differences between the means of the two groups were calculated by two-tailed Student’s t-test with nonparametric Mann-Whitney tests. Significant statistical difference was expressed as follows - **p < 0.01, ***p < 0.001; n=5-7.

### Non-iNKT cells significantly differ from the iNKTs both quantitatively and qualitatively between the liver and spleen of WT mice

In addition to the quantity of the cells, we also determined the phenotype of these cells by analyzing the expression of the co-receptor molecules CD4 and CD8, the activation marker CD69, and the intracellular cytokines IFN-γ and IL-4. We found that the non-iNKTs are either double negative (DN) or CD8^+^ in the liver and spleen. In contrast, the iNKTs are mostly CD4^+^ with a lesser extent of DNs (Fig. 2). Furthermore, at baseline, iNKTs are significantly more CD69^+^ compared to the non-iNKTs in both organs (49.13±4.92 vs 17.61±2.18) (Fig. 2). Number of intracellular IFN-γ positive iNKTs are significantly more than non-iNKTs in the liver (6.21±0.7 vs 3.428±0.66), but this expression of IFN-γ reversed to the opposite direction in the spleen (4.522±0.56 vs 11.74±1.51) (Fig. 2). Conversely, the IL-4^+^ non-iNKTs are significantly more in liver and spleen compared to the IL-4^+^ iNKTs (4.448±1.27 vs 2.006±0.3 and 12.88±1.78 vs 5.848±1.14) (Fig. 2). This quantitative and qualitative analysis distinguishes discretely the iNKT and non-NKT subsets in the liver and spleen.

**Figure. 2.**
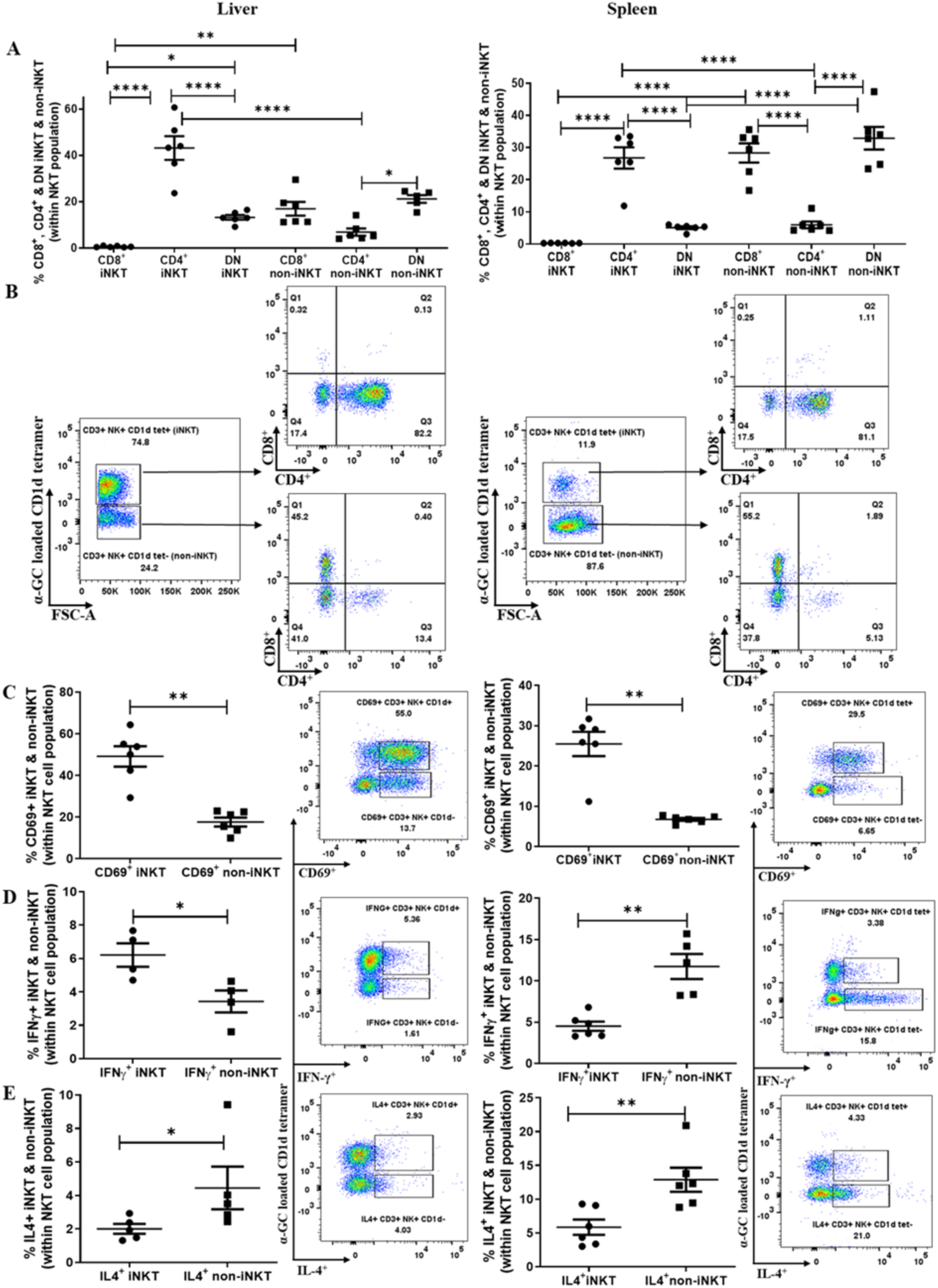
Qualitative analysis of non-iNKT cells in the liver and spleen of WT mice. (A) Scatter plots denoting the number of iNKT and the non-iNKTs expressing co-receptor markers CD4, CD8 and CD4-CD8-(DN) within the NKTs population in liver and spleen. (B) Representative dot plots showing the CD4, CD8 and DN gates within the non-iNKT and iNKT gates in liver and spleen. The gates - Q1 represent CD8^+^, Q3 is CD4^+^ and Q4 is DN population. (C) Scatter plot denoting the expression of activation marker CD69^+^ iNKT and non-iNKTs within the NKTs, followed by representative dot plots in the liver and spleen. (D) Scatter plot denoting the intracellular IFN-γ^+^ iNKT and non-iNKTs within the NKTs, followed by representative dot plots in the liver and spleen. (E) Scatter plot denoting the intracellular IL-4^+^ iNKT and non-iNKTs within the NKTs, followed by representative dot plots in the liver and spleen. All data are represented as Mean ± SEM. Statistical difference between the means of two groups are calculated by two-tailed Student’s t-test with nonparametric Mann-Whitney tests. For evaluating the difference between multiple group one-way analysis of variance (ANOVA) with Dunn’s multiple comparison post-test was performed. Significant statistical difference was expressed as follows - *p < 0.05, **p < 0.01, ***p < 0.001, and ****p < 0.0001, ns - non-significant.

### Lack of Apoe gene increases non-iNKT numbers irrespective of diet-induced atherosclerosis

To determine the role of the non-iNKT cells in diet-induced atherosclerosis, we investigated these cells in HFD-fed atheroprone *Apoe^-/-^* mice. Atherosclerosis development was confirmed by an increased lipid profile from the sera and a significant increase in the lesion area in the HFD-fed *Apoe^-/-^* compared to the chow group (Fig. 3). However, there was no significant change in the body weight, liver enzymes, and the weight and size of the liver and spleen of the chow and HFD fed *Apoe^-/-^* mice over the 20 weeks (Fig. 3).

**Figure. 3.**
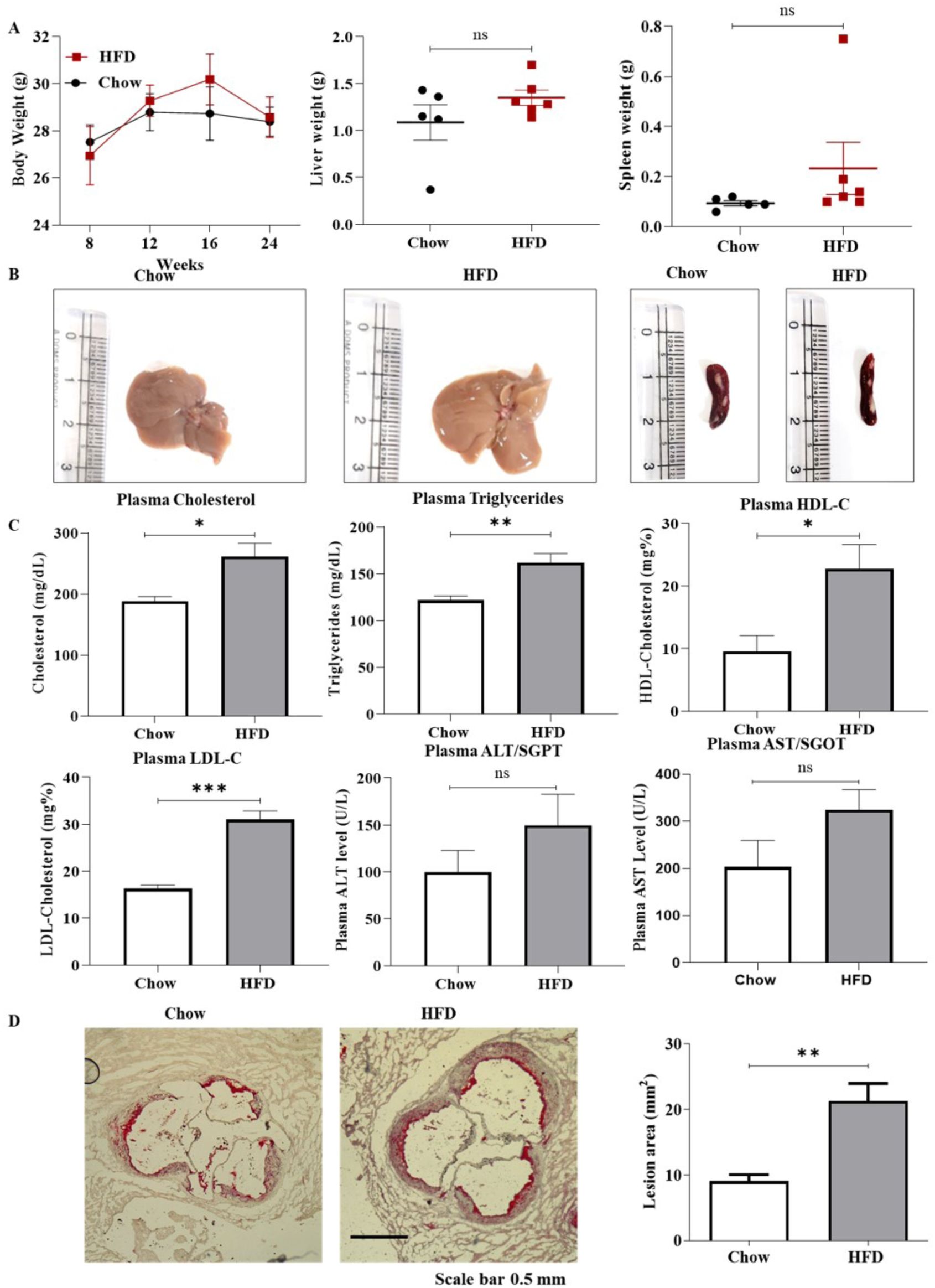
Atherosclerosis development in HFD fed *Apoe^-/-^* mice compared to chow fed *Apoe^-/-^* mice. (A) Body weight of both the groups after starting HFD at 8, 12, 16 and 24 weeks of the age of the mice (n=12-15). HFD started from 8 weeks. Liver and spleen weight of the two groups after sacrificing the mice. (B) Representative images of the liver and spleen from chow and HFD fed mice depicting no change in their size. (C) Bar plots representing the plasma levels of cholesterol, triglycerides, HDL-C, LDL-C, ALT and AST in the chow and HFD fed mice (n=7-8). (D) Oil-red O staining of the aortic valves in the chow and HFD, denoting lipid deposition and the bar plots represent the lesion area in mm^2^. Scale Bar – 0.5mm. All data are represented as Mean ± SEM. Statistical difference between the means of two groups are calculated by two-tailed Student’s t-test with nonparametric Mann-Whitney tests. Significant statistical difference was expressed as follows - \**p* < 0.05, \*\**p* < 0.01, \*\*\**p* < 0.001, and \*\*\*\**p* < 0.0001, ns - non-significant.

Our data did not reveal any significant change in the whole NKT cell population between the chow and HFD groups of *Apoe^-/-^* (Fig. 4). We did not find any significant change in the non-iNKT numbers in the liver and spleen between the chow and HFD groups (l69.39±4.23% vs 70.68±3.93%; 91.84±1.65 vs 92.33±2.55, respectively). The number of the iNKT subsets also remain unchanged between the chow and HFD groups. However, the non-iNKTs were always significantly increased compared to the iNKTs in both the liver (Chow – 69.39±4.23% vs 31.20±3.54%; HFD – 70.68±3.94% vs 30.58±3.13%) and spleen (Chow – 91.84±1.65% vs 7.091±1.54%; HFD – 92.33±2.55% vs 4.582±1.47%) of chow and HFD-fed *Apoe* null mice. The increase of the non-iNKTs compared to the iNKTs were also observed in the HFD-fed aorta (68.4±5.36 vs 29±5.41) (Fig. 4).

**Figure. 4.**
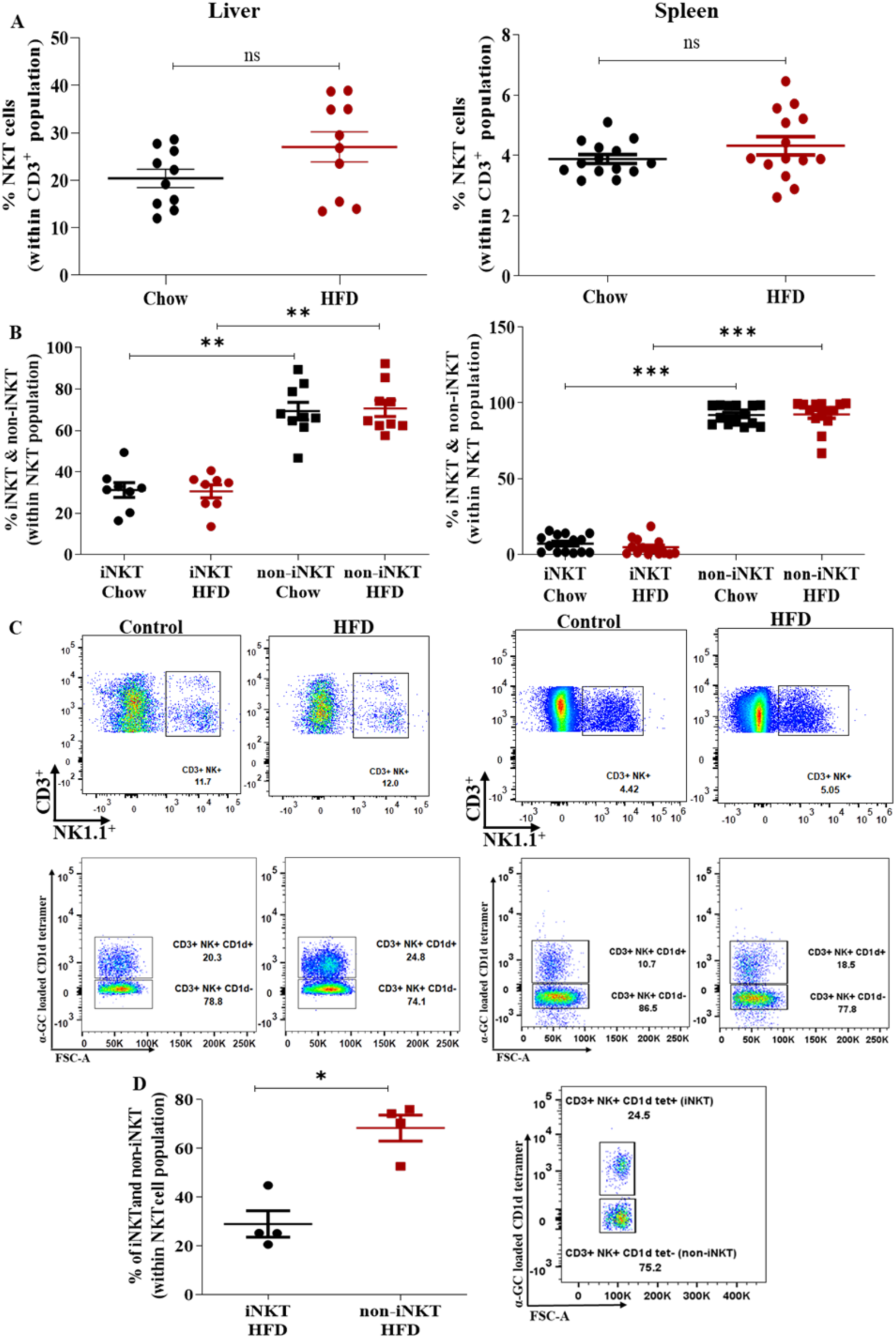
Quantitative analysis of non-iNKT cells in the liver and spleen of chow and HFD fed *Apoe^-/-^* mice. (A) Scatter plots denoting the number of NKT cells within the CD3+ T cell population in liver and spleen of chow and HFD *fed Apoe^-/-^* mice. (B) Scatter plots denoting the number of the iNKT and the non-iNKTs within the NKT population in chow and HFD fed liver and spleen of *Apoe^-/-^* mice. (C) Representative dot plots showing the gates of NKT and the non-iNKT and the iNKT population in the liver and spleen of chow and HFD fed *Apoe^-/-^* mice. (D) Number of the iNKT and non-iNKTs within the NKT population isolated from the plaques deposited in the aorta of the HFD fed *Apoe^-/-^* mice. All data are represented as Mean ± SEM. Statistical difference between the means of two groups are calculated by two-tailed Student’s t- test with nonparametric Mann-Whitney tests. For evaluating the difference between multiple group one-way analysis of variance (ANOVA) with Dunn’s multiple comparison post-test was performed. Significant statistical difference was expressed as follows - *p < 0.05, **p < 0.01, ***p < 0.001, and ****p < 0.0001, ns - non-significant.

### Apoe gene reduces the overall non-iNKT numbers and the activated CD69^+^ non-iNKTs but increases the IL-4^+^ non-iNKTs

The previous data revealed no significant difference in the numbers of the non-iNKT cells in the chow and HFD-fed *Apoe^-/-^* mice; however, we observed an inherent increase of the non-iNKTs compared to the iNKTs irrespective of atherosclerosis. These data arise the doubt on the absence of the *Apoe* gene in these mice that might affect the cell numbers. Accordingly, we were inquisitive to discern the baseline differences in the non-iNKTs quantitatively and qualitatively in the cells with and without ApoE. Therefore, we used WT and *Apoe^-/-^* mice that were 8-10 weeks of age and were provided with standard chow only (Fig. 5). From our data, we found unanticipated results, that the whole NKT population was increased in the liver by 32.5% but decreased in the spleen by 30.3% in *Apoe^-/-^* compared to the WT (Fig. 5). These data revealed a significant increase in the non-iNKT numbers in the *Apoe^-/-^* mice by 87.5% in the liver and 43.8% in the spleen compared to the WT controls (Fig. 5). Whereas the iNKT numbers reduced significantly in the *Apoe^-/-^* mice compared to the WT. In the qualitative analysis, we found that the CD8^+^ non-iNKTs are significantly increased, almost doubled in the liver of *Apoe* null mice compared to the WT (34.9±2.39 vs 17.15±2.87) (Fig. 6). The CD4^+^ non-iNKTs decreased significantly in the liver of the *Apoe^-/-^* (2.659±0.21 vs 6.903±1.59) (Fig. 6). The CD4^+^ iNKT numbers also significantly reduced in the liver and spleen of the *Apoe^-/-^* mice compared to the WT controls (Fig. 6). The DN non-iNKTs are also increased both in the liver and spleen (37.08±2.15 vs 19.31±2.32 and 62.78±3.19 vs 32.9±3.52) of the *Apoe^-/-^* compared to the WT depicting the effect of *Apoe* gene on these cells (Fig. 6). The CD69 activation status and the cells positive for this marker are significantly increased in the *Apoe^-/-^* in both liver and spleen (54.6±2.8 vs 17.61±2.18 and 22.79±3.54 vs 6.79±0.365) whereas CD69^+^ iNKTs decrease significantly in the liver and spleen of the *Apoe^-/-^* compared to the WT (Fig. 7). However, there was no difference in the numbers of the IFN-γ^+^ non-iNKT cells in between these two mice strains but the IFN-γ^+^ iNKTs decreased significantly in the *Apoe^-/-^* mice. It is noteworthy that the IFN-γ^+^ non-iNKTs were significantly increased by 86.2% compared to IFN- γ^+^ iNKTs in the liver of the *Apoe^-/-^* mice but not in the WT mice. However, in the spleen of both the WT and *Apoe^-/-^* mice the IFN-γ^+^ non-iNKTs were significantly increased than the IFN- γ^+^ iNKTs (Fig. 7). IL-4 expressing non-iNKTs increased significantly in both the liver and spleen (3.518±0.43 vs 1.16±0.18 and 11.27±0.93 vs 1.62±0.26) of WT compared to the *Apoe^-/-^* mice (Fig. 7).

**Figure. 5.**
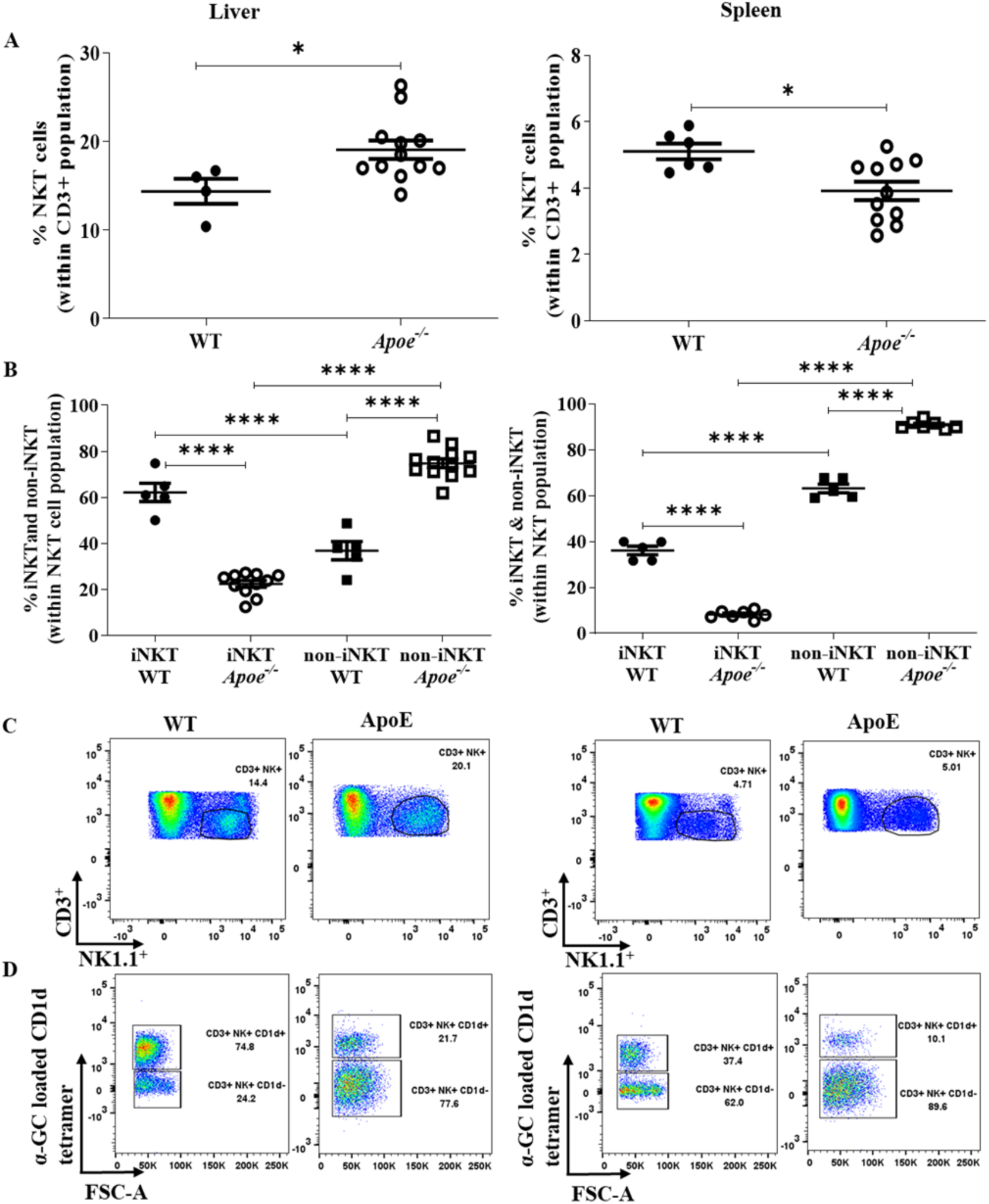
Baseline quantitative analysis of NKT, non-iNKT and iNKT cells in the liver and spleen of 8-10 weeks old chow fed WT and *Apoe^-/-^* mice. (A) Scatter plots denoting the number of NKT cells within the CD3^+^ T cell population in liver and spleen of chow fed WT and *Apoe^-/-^* mice. (B) Scatter plots denoting the number of the iNKTs and the non-iNKTs within the NKT population in liver and spleen of chow fed WT and *Apoe^-/-^* mice. (C), (D) Representative dot plots showing the gates of NKT population and the non-iNKT and the iNKT population in the liver and spleen respectively of WT and *Apoe^-/-^* mice. All data are represented as Mean ± SEM. Statistical difference between the means of two groups are calculated by two-tailed Student’s t-test with nonparametric Mann-Whitney tests. For evaluating the difference between multiple group one-way analysis of variance (ANOVA) with Dunn’s multiple comparison post-test was performed. Significant statistical difference was expressed as follows - *p < 0.05, **p < 0.01, ***p < 0.001, and ****p < 0.0001, ns - non-significant.

**Figure. 6.**
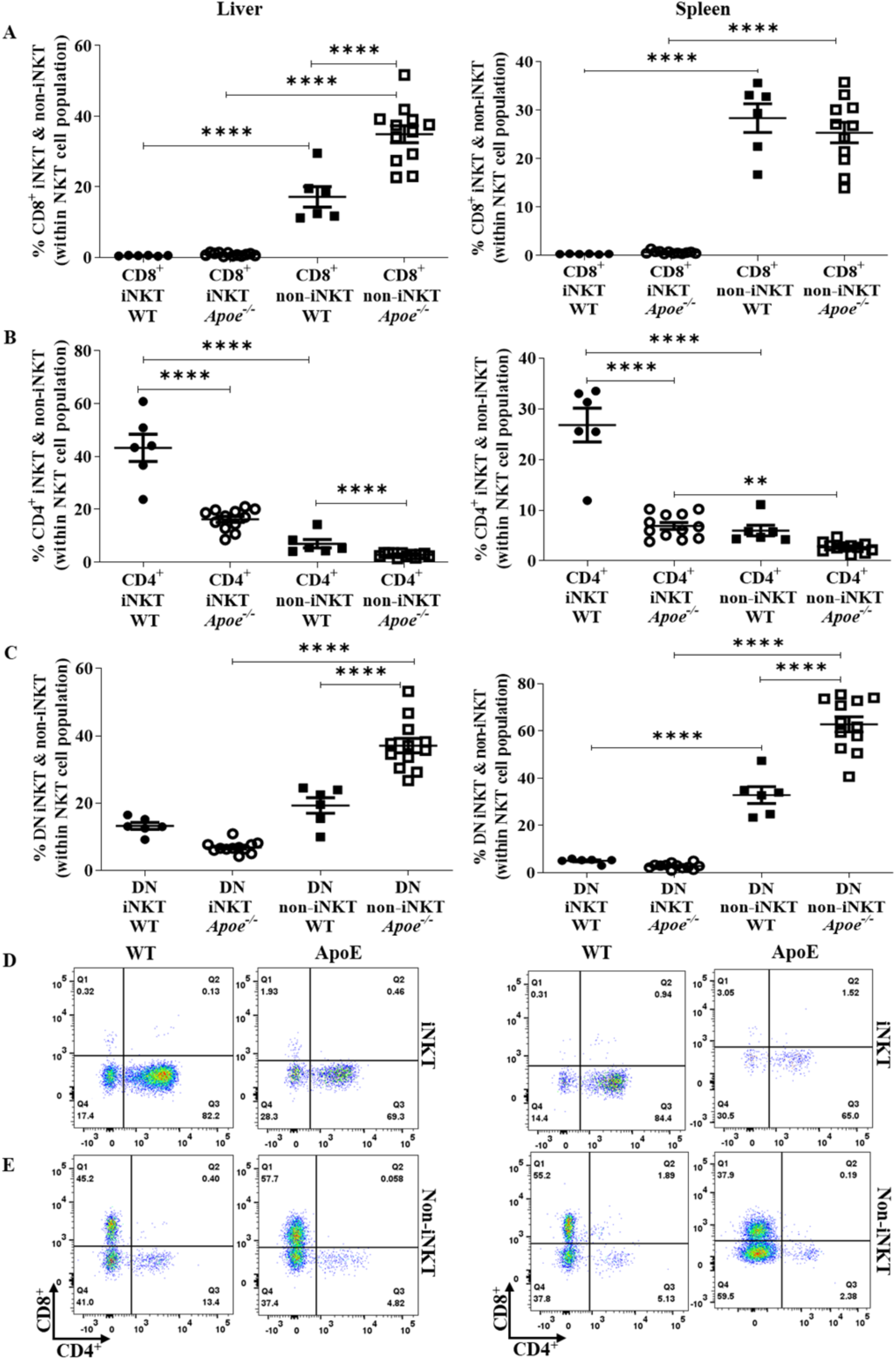
Baseline co-receptor expression differences in the iNKTs and the non-iNKTs within the NKT cell population in the liver and spleen of 8-10 weeks old chow fed WT and *Apoe^-/-^* mice. (A), (B), (C) Scatter plots denoting the number of the iNKTs and non-iNKTs expressing CD8^+^, CD4^+^, and CD8^-^CD4^-^ (DN) within the NKT population in the liver and spleen of chow fed WT and *Apoe^-/-^* mice. (D), (E) Representative dot plots showing the gates of non-iNKTs and the iNKTs expressing CD8^+^, CD4^+^, and DN within the NKTs in the liver and spleen of chow fed WT and *Apoe^-/-^* mice. All data are represented as Mean ± SEM. For evaluating the difference between multiple group one-way analysis of variance (ANOVA) with Dunn’s multiple comparison post-test was performed. Significant statistical difference was expressed as follows - \**p* < 0.05, \*\**p* < 0.01, \*\*\**p* < 0.001, and \*\*\*\**p* < 0.0001, ns - non-significant.

**Figure. 7.**
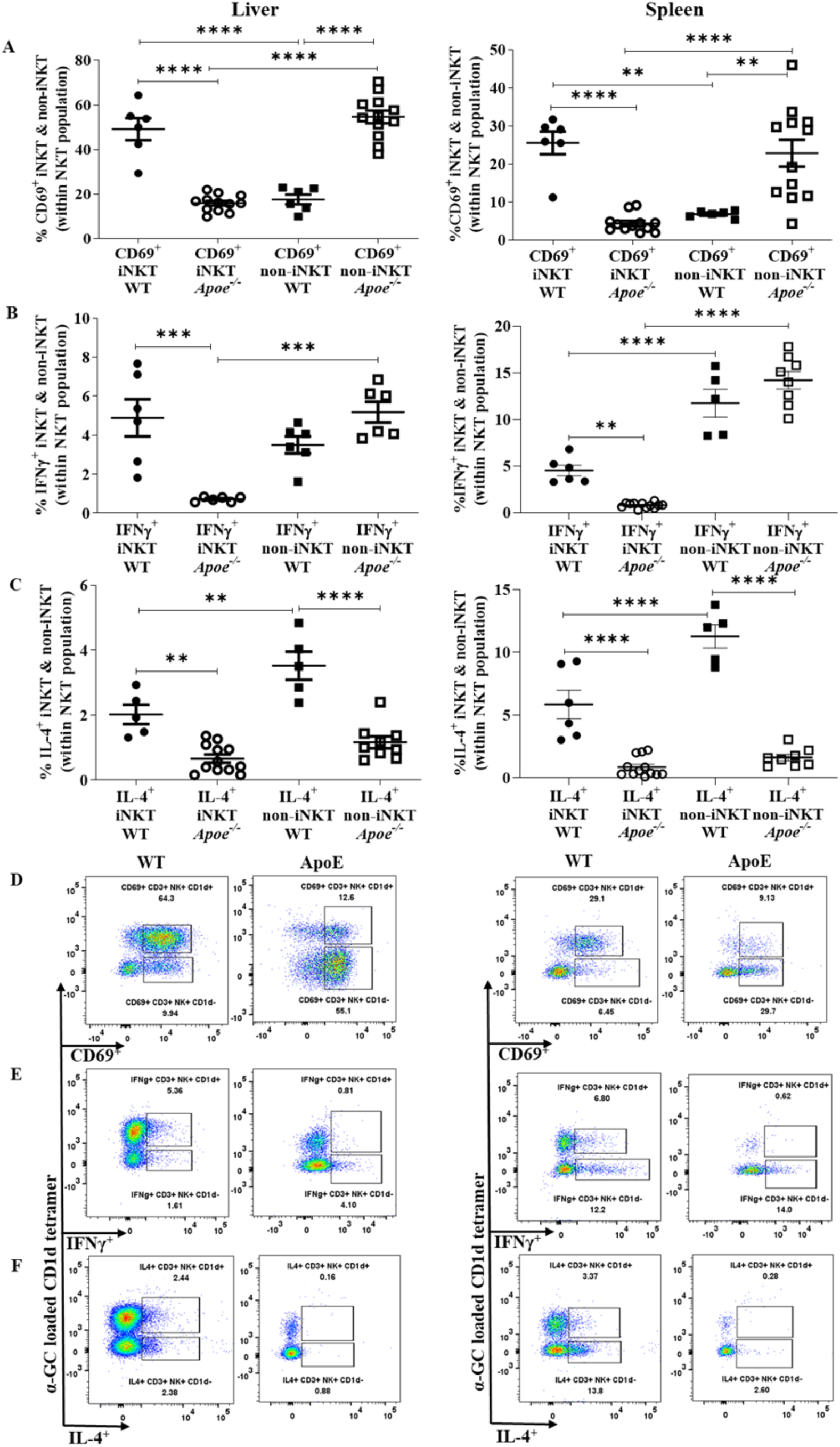
Baseline analysis of activation marker and intracellular cytokine expression of iNKT and non-iNKT cells in the liver and spleen of 8-10 weeks old chow fed WT and *Apoe^-/-^* mice. (A), (B), (C) Scatter plots denoting the number of the CD69^+^, IFNγ^+^ and IL-4^+^ iNKT and the non-iNKTs within the NKT population in liver and spleen of WT and *Apoe^-/-^* mice respectively. (D), (E), (F) Representative dot plots showing the CD69^+^, IFNγ^+^ and IL-4^+^ non-iNKT and the iNKTs. All data are represented as Mean ± SEM. For evaluating the difference between multiple group one-way analysis of variance (ANOVA) with Dunn’s multiple comparison post-test was performed. Significant statistical difference was expressed as follows - \**p* < 0.05, \*\**p* < 0.01, \*\*\**p* < 0.001, and \*\*\*\**p* < 0.0001, ns - non-significant.

### Non-iNKTs play important immune-regulatory roles in dyslipidemia of WT

It is known that WT mice are resilient in developing diet-induced atherosclerosis, irrespective of the duration of HFD feeding. However, dyslipidemia developed in these WT mice after feeding them HFD for a similar duration as fed to *Apoe^-/-^* mice (20 weeks) (Fig. 8). Plasma cholesterol levels and LDL-C levels increased significantly in the HFD-fed group, while triglycerides, HDL-C, and liver enzymes and body weight were unchanged without any lipid deposition in the aortic valves (Fig. 8). The number of the non-iNKTs in the liver was significantly increased by 46% in the HFD-fed WT compared to chow fed group, contrastingly, the non-iNKTs in the spleen were unchanged (Fig. 9). The CD8^+^ non-iNKTs were significantly decreased by 68.6% in the liver and doubly increased in the spleen of the HFD-fed group compared to the chow-fed WT (Fig. 10). The CD4^+^ non-iNKTs decreased more than two times in the HFD-fed WT liver whereas the CD4+ iNKTs decreased by 53.7% in the HFD-fed spleen (Fig. 10). In the spleen, the DN non-iNKTs also increased by 39.3%, following the similar trend of CD8^+^ non-iNKTs (Fig. 10). The CD69^+^ iNKTs decreased in the HFD-fed spleen by 44.9% and the CD69^+^ non-iNKTs were significantly less than the CD69^+^ iNKTs in both the liver and spleen of chow and HFD groups (Fig. 11). The intracellular IFN-γ^+^ non-iNKT cells increased twice in their numbers in the HFD-fed spleen compared to the chow-fed WT (Fig. 11). No significant change was observed in the activation status and intracellular IL-4 expression of the non-iNKTs between the chow and HFD fed group in WT mice (Fig. 11). Therefore, from these data, it is evident that the non-iNKTs are involved in the immune response in dyslipidemic conditions. On the contrary, these differences in the numbers of non-iNKTs observed in diet-induced dyslipidemic WT were not found in the diet-induced atherosclerotic *Apoe^-/-^* mice.

**Figure. 8.**
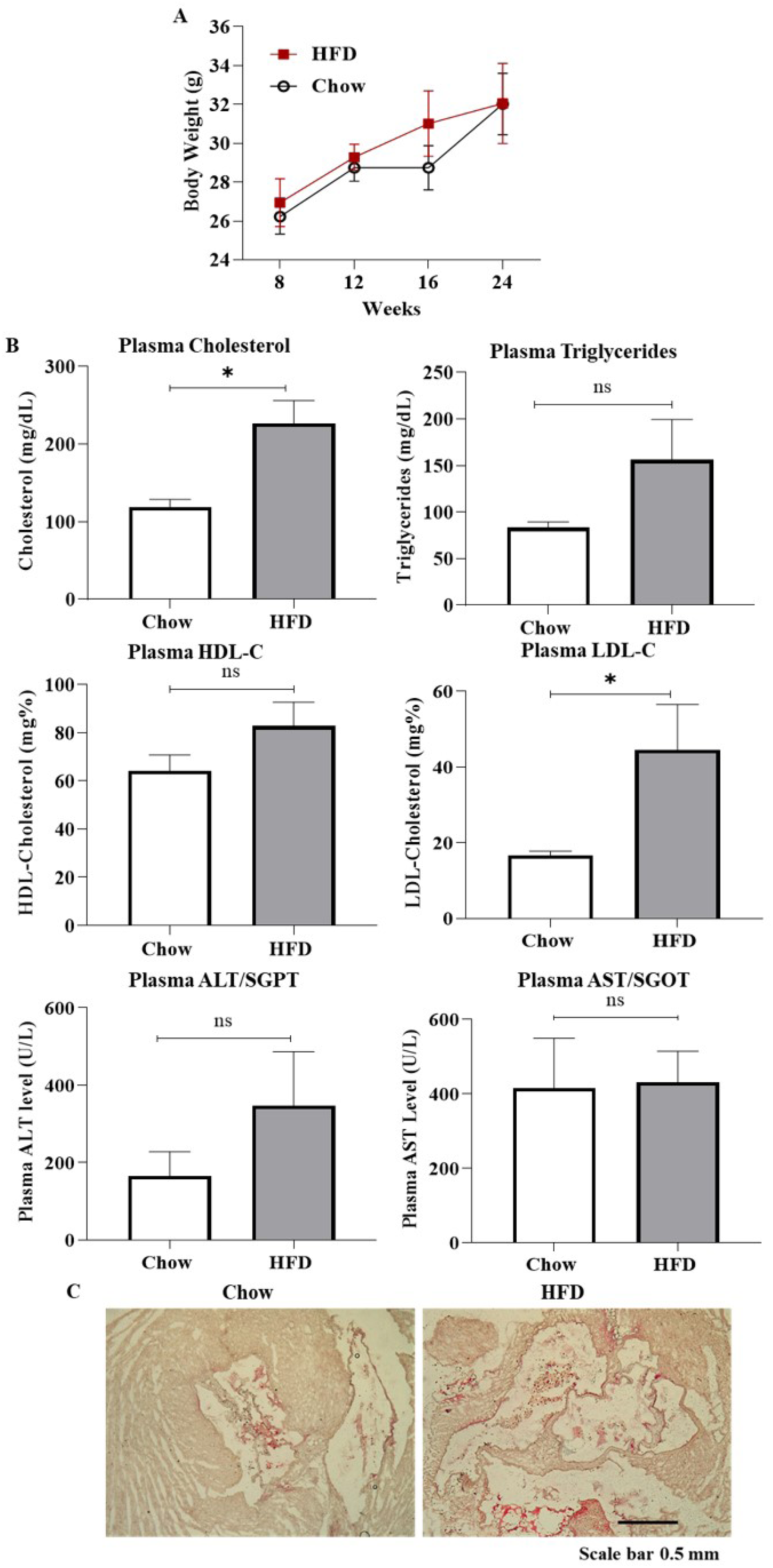
Development of dyslipidemia in HFD fed WT compared to chow fed WT mice. (A) Body weight of both the groups after starting HFD at regular intervals (n=10-14). (B) Bar plots representing the plasma levels of cholesterol, triglycerides, HDL-C, LDL-C and liver enzymes, ALT and AST in the chow and HFD fed mice (n=7-8). (C) Oil-red O staining of the aortic valves in the chow and HFD, denoting no lipid deposition in WT. Scale Bar – 0.5mm. All data are represented as Mean ± SEM. Statistical difference between the means of two groups are calculated by two-tailed Student’s t-test with nonparametric Mann-Whitney tests. Significant statistical difference was expressed as follows - \**p* < 0.05, ns - non-significant.

**Figure. 9.**
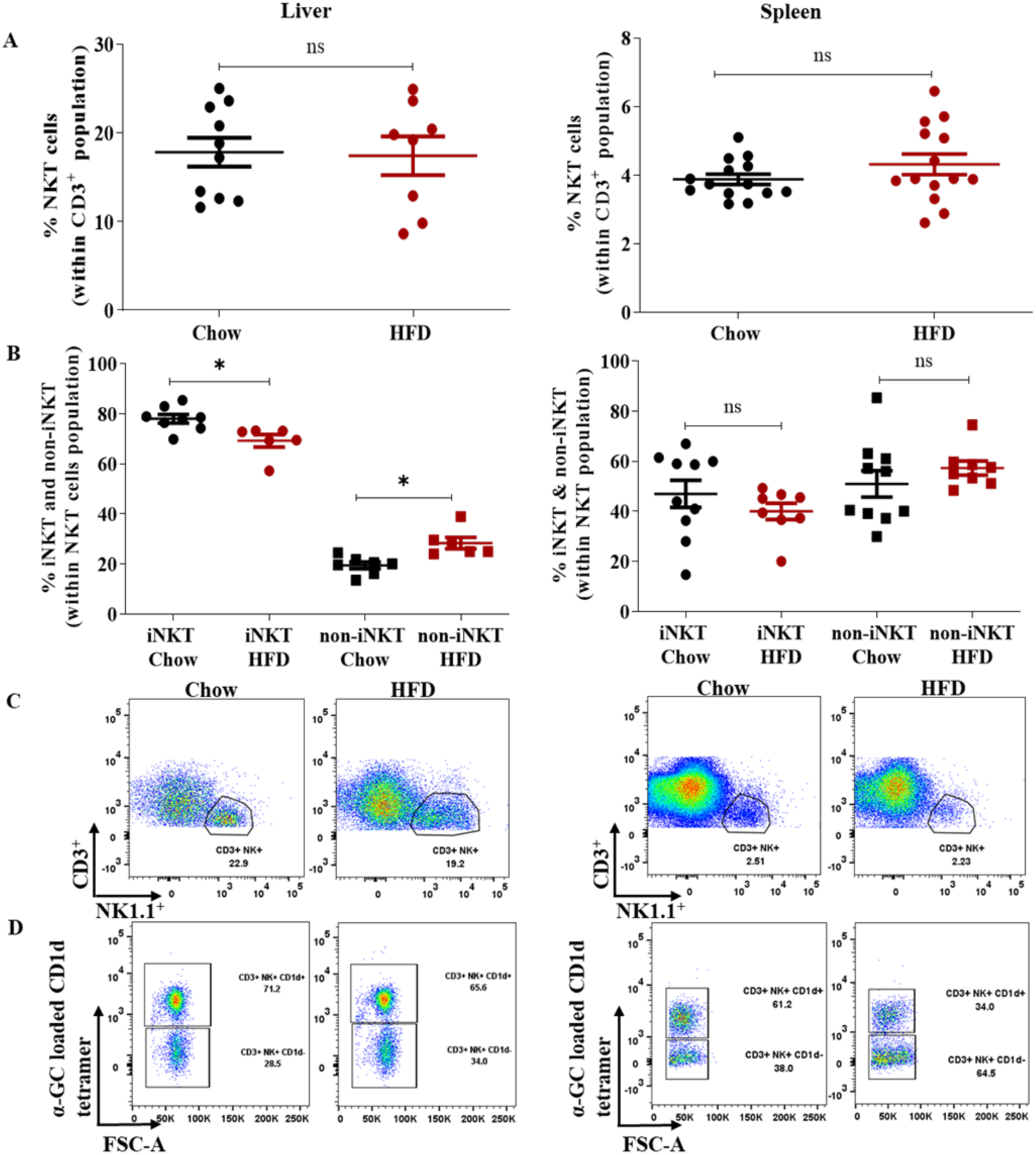
Quantitative analysis of NKT, non-iNKT and iNKT cells in the liver and spleen of chow and HFD fed WT mice. (A) Scatter plots representing the number of NKT cells within the CD3^+^ T cell population in liver and spleen of chow and HFD fed WT. (B) Scatter plots representing the number of the iNKT and the non-iNKTs within the NKT population in liver and spleen of chow and HFD fed WT. (C), (D) Representative dot plots showing the gates of NKT population and the non-iNKT and the iNKT population in the liver and spleen respectively of chow and HFD fed WT. All data are represented as Mean ± SEM. Statistical difference between the means of two groups are calculated by two-tailed Student’s t-test with nonparametric Mann-Whitney tests. For evaluating, the difference between multiple group one-way analysis of variance (ANOVA) with Dunn’s multiple comparison post-test was performed. Significant statistical difference was expressed as follows - *p < 0.05, ns - non-significant.

**Figure. 10.**
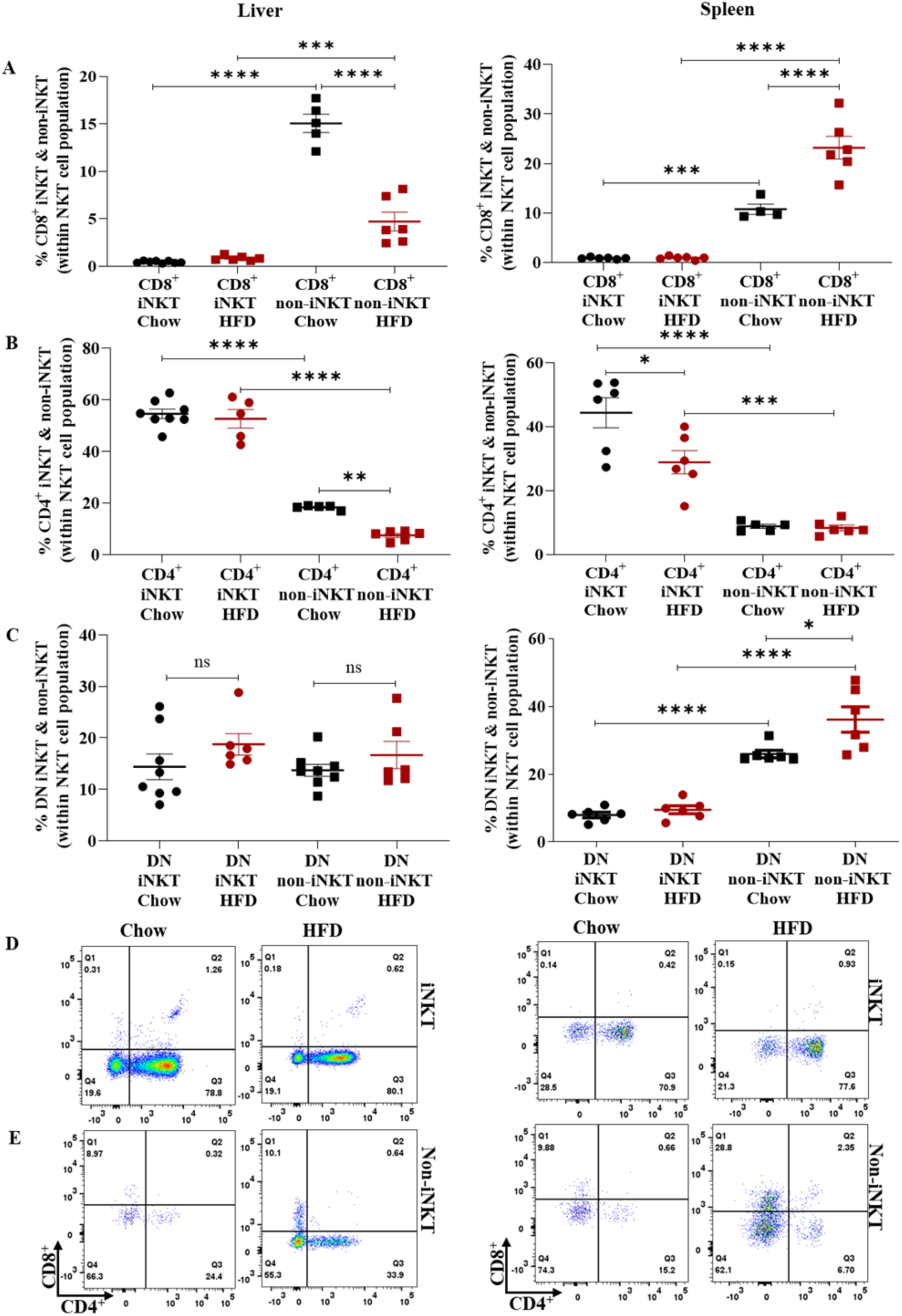
Qualitative analysis of the co-receptor expression differences in the non-iNKTs and the iNKTs within the NKT cell population in the liver and spleen of chow and HFD fed WT mice. (A), (B), (C) Scatter plots representing the number of the iNKTs and the non-iNKTs expressing CD8^+^, CD4^+^, and CD8^-^CD4^-^ (DN) within the NKT population in the liver and spleen of chow and HFD fed WT mice. (D), (E) Representative dot plots showing the gates of non-iNKTs and the iNKTs expressing CD8^+^, CD4^+^, and DN within the NKTs in the liver and spleen of chow and HFD fed WT mice. All data are represented as Mean ± SEM. For evaluating the difference between multiple group one-way analysis of variance (ANOVA) with Dunn’s multiple comparison post-test was performed. Significant statistical difference was expressed as follows - \**p* < 0.05, \*\**p* < 0.01, \*\*\**p* < 0.001, and \*\*\*\**p* < 0.0001, ns - non-significant.

**Figure. 11.**
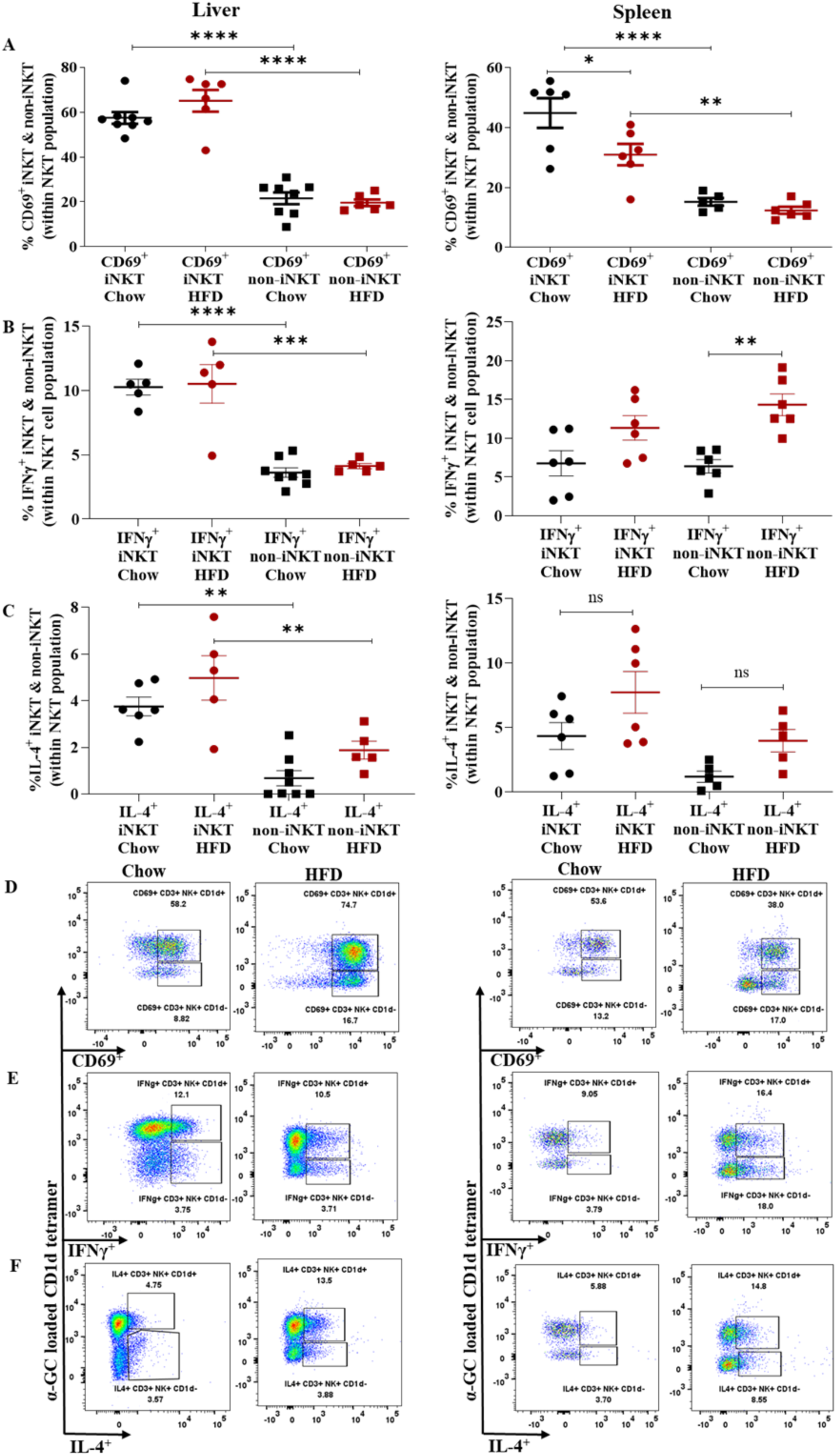
Qualitative analysis of activation marker and intracellular cytokine expression of non-iNKT cells in the liver and spleen of chow and HFD fed WT mice. (A), (B), (C) Scatter plots representing the number of the CD69^+^, IFNγ^+,^ and IL- 4^+^ iNKT and the non-iNKTs within the NKT population in liver and spleen of chow and HFD fed WT mice respectively. (D), (E), (F) Representative dot plots showing the CD69^+^, IFNγ^+,^ and IL-4^+^ non-iNKT and the iNKTs. All data are represented as Mean ± SEM. For evaluating the difference between multiple group one-way analysis of variance (ANOVA) with Dunn’s multiple comparison post-test was performed. Significant statistical difference was expressed as follows - \**p* < 0.05, \*\**p* < 0.01, \*\*\**p* < 0.001, and \*\*\*\**p* < 0.0001, ns - non-significant.

## Discussion

In this study, we developed a method of non-iNKT cell identification in mice using a negative selection strategy with the help of flow cytometry. The non-iNKT cells are less than the iNKTs in the liver contrary to the spleen, where the non-iNKTs are more than the iNKTs. This difference in the frequency of the NKT subsets between the homing organ liver and the lymphoid organ spleen marks their immune-regulatory function to maintain homeostasis. Their distinct qualitative features follow the quantitative differences as well. We analyzed these cells based on co-receptor expression, such as CD4, CD8, and CD4^-^CD8^-^ (DN), activation marker expression, CD69, and intracellular cytokines IFN-γ and IL-4.

Our data reveals that the non-iNKT cells are predominantly DN and/or CD8^+^, but their activation status is lower than the iNKTs. However, we found an increased number of IL-4^+^ non-iNKTs and reduced IFN-γ^+^ non-iNKTs compared to the iNKT subset in the liver. Moreover, in the spleen, intracellular expression of both the cytokines IFN-γ and IL-4 increased in the non-iNKT subset compared to the iNKTs. Also, from the previous reports, it is evident that iNKT cells respond to rapid antigenic stimulation by production of both Th1 cytokines (IFN-γ and TNF-α) and Th2 cytokines (IL-4, IL-10, IL-13) ^34, 35^. Further, our data indicates that the non-iNKTs are poised for Th2 response in the homing organ liver due to increased IL-4. Additionally, the non-iNKTs that localize in the spleen exhibit Th1 and Th2 response due to higher levels of IFN-γ and IL-4 than iNKTs. Our data here takes the previous knowledge one step further and suggests that, although the activated CD69^+^ iNKTs are higher compared to the non-iNKTs, yet the non-iNKTs are poised for increased Th2 response in the liver and both Th1 and Th2 response in the spleen.

*Apoe* null mice on HFD develop atherosclerosis and are considered a bonafide model for studying atherosclerosis. In our experiments, despite HFD-induced atherosclerosis in the *Apoe^-/-^* mice, we did not find any change in the quantity of these cells in the liver and spleen, indicating a systemic increase of the non-iNKTs in the *Apoe^-/-^* mice irrespective of the atherosclerosis development. Furthermore, we analyzed the immune cells from the plaques developed in the aorta of only the HFD-fed *Apoe^-/-^* mice, and results were similar to the liver and spleen, i.e., the non-iNKT cells are increased compared to the iNKTs. It is noteworthy to mention that the atherosclerotic plaque develops only in the aorta of the HFD-fed *Apoe^-/-^* mice and not in chow-fed *Apoe^-/-^* mice. Our data shows an increase of non-iNKTs in the atherosclerotic plaques of the aorta of the HFD-fed *Apoe^-/-^* mice, which corroborates with our previous data of the systemic increase of non-iNKTs in the liver and spleen of the *Apoe^-/-^* mice irrespective of diet. Nonetheless, this is the first study to report the non-iNKTs in the atherosclerosis of *Apoe^-/-^* mice. This data was unanticipated since atherosclerosis is a hyperlipidemic disease, and *Apoe^-/-^* mice are a gold standard model for studying the pathogenesis of this disease. Furthermore, the NKT cells recognize glycolipids presented by ubiquitously expressed CD1d molecules that are non-classical Major Histocompatibility Complex (MHC) Class-I.

To better understand the probable reasons, we investigated the differences in the non-iNKT cell quantity and quality between the young, chow-fed WT and *Apoe^-/-^* mice strains. These data concomitantly revealed that the absence of the *Apoe* gene leads to an increase in the number of non-iNKTs in *Apoe^-/-^* mice. The data shows that the presence of the *Apoe* gene in the WT mice leads to the variation in the numbers and phenotype of the NKT subsets. The non-iNKT numbers significantly increased, whereas the iNKTs significantly decreased in the *Apoe* null mice. The decrease of the iNKT numbers was also reported by Braun *et al*. ^36^. This group reported that the iNKT cell numbers decreased in the liver, spleen, and thymus of the *Apoe^-/-^* mice compared to WT. Furthermore, their data reveals similar expression of CD1d on the splenic DCs, indicating appropriate antigen-presentation in both strains. However, co-culturing of the splenic DCs from WT and *Apoe^-/-^* with iNKTs from WT and *Apoe^-/-^* exhibited non-functional iNKTs ^36^. They defined this non-functionality of the iNKTs in the *Apoe^-/-^* as “spontaneous anergy and hyporesponsiveness”. Therefore, our data corroborating with this report takes the understanding of the NKT population one step further and indicates that the inherent increase of the non-iNKTs might be a compensation mechanism of the system due to decreased iNKTs in the *Apoe^-/-^* mice. Although there are reports about the importance of *Apoe-*mediated delivery of the lipids to the endosomal compartment of the antigen-presenting cells (APCs) for presentation by the CD1 molecule, on the contrary, the report from Braun *et al*. provides vital evidence about the proper functioning of the CD1d molecule in the *Apoe^-/-^* mice and different functioning of the iNKT subset ^37^. Therefore, it is evident that the absence of *Apoe* does not hinder the presentation of glycolipids through CD1d molecule in the APCs; however, ApoE is inevitable in maintaining proper numbers of the NKT subsets. Further, in the liver of the *Apoe^-/-^* mice, the CD8^+^ and the DN non-iNKTs are increased, whereas the CD4^+^ non-iNKTs and the CD4^+^ iNKTs are decreased. The CD4^+^ iNKTs also decrease in the spleen of the *Apoe^-/-^* mice. Therefore, the iNKT and the non-iNKT cells expressing the CD4 co-receptor are suppressed without ApoE. Furthermore, the co-receptor expressing non-iNKT subpopulation (CD8^+^, CD4^+^, and DN) is affected in the liver of the *Apoe* null mice; this exhibits the critical mechanisms of the *Apoe* gene on these cells. The scope of the present study is limited only to the observation of the effect of the ApoE on the non-iNKT subset without delving into the pathways of how this gene is affecting the subsets of the NKT population. Nevertheless, our data reveals for the first time that the *Apoe* gene affects the non-iNKT subset and, therefore, warrants using the *Apoe^-/-^* mice to study the non-iNKTs. ApoE also affects the activation of the non-iNKTs. The CD69^+^ non-iNKTs significantly increase in the liver and spleen of the *Apoe^-/-^* mice. Our data corroborates with the results published by Braun *et al.,* which state that the CD69^+^ iNKTs significantly decrease in the liver and spleen of the *Apoe^-/-^* mice compared to the WT. The excessive increase in the CD69^+^ non-iNKTs can be a compensatory mechanism of the system due to the subdued CD69^+^ iNKTs. However, the IFN-γ^+^ non-iNKTs are increased significantly than the IFN-γ^+^ iNKTs in the liver of the *Apoe^-/-^* mice, unlike the WT populations. This data delineates that the absence of ApoE renders a pro-inflammatory phenotype to the non-iNKTs in their homing organ liver. Moreover, the IL-4^+^ non-iNKTs significantly decrease in the *Apoe^-/-^* mice, indicating a suppressed Th2 response. Our work contributes substantial confirmation and advances toward understanding the *Apoe* gene’s effects on the non-iNKT subset, which were previously unknown; nonetheless, how this gene affects the non-iNKTs still needs to be further investigated.

The glycolipid antigens activate NKT cells, and dyslipidemia is the primary cause of atherosclerosis. Subsequently, HFD-fed WT mice developed dyslipidemia that manifested with significantly high cholesterol and LDL-C in plasma. We used this dyslipidemic WT mice model to determine the effect of a deranged lipids’ milieu on the biology of non-iNKTs. The results delineated increased non-iNKT cell numbers in the liver but not in the spleen of the HFD WT group compared to chow, exhibiting that these cells are homed in large numbers in the liver as a response to augmented lipids in the milieu. On the other hand, corroborating with previously published reports from other groups, the iNKT cell numbers significantly reduced in the HFD-fed WT, probably due to the continuous stimulation from the increased lipids resulting in the occurrence of anergy in these cells ^9, 16, 17^. This also correlates with the higher activation status of the CD69^+^ iNKT compared to the CD69^+^ non-iNKTs in the baseline (Supplementary Fig. S3A). Therefore, it might be possible that to counter-balance the reduced iNKTs, the frequency of the non-iNKT subset increases in the liver of the HFD-fed WT mice, similar to the phenotype exhibited by the *Apoe* null mice. Furthermore, our results also demonstrate that both the CD8^+^ and the DN non-iNKTs significantly increase in the spleen of the HFD group, whereas, in the liver only the CD8^+^ non-iNKTs increase. Additionally, proinflammatory IFN-γ^+^ non-iNKTs are aggravated in the spleen of the HFD group in WT. Hence, this amelioration of the CD8^+^, DN, and the IFN-γ^+^ non-iNKTs in the spleen of the HFD-fed WT mice demonstrates the participation of these cells in regulating the immune response due to the inflammatory conditions caused by dyslipidemia. However, the cellular mechanism of action is yet to be deciphered. Consequently, these data imply the crucial effect of the presence of the *Apoe* gene in the WT that leads to the regular functioning of the non-iNKT subsets, unlike the disrupted functionality of these cells in *Apoe^-/-^* mice.

There are proceedings on the research related to the potential role of iNKTs in experimental murine models of cardiovascular diseases that might pioneer the applications for human CVDs in the future ^38^. Nonetheless, our work alters the current understanding of NKT cell biology and warrants using the *Apoe^-/-^* mice to investigate the NKT population. This study establishes the fact that the non-iNKT subset gets affected in the *Apoe* null mice due to the ablated *Apoe* gene, which is otherwise functional and exhibits immune regulatory functions in dyslipidemia of WT mice strain; however, the cellular mechanism of these observations are yet to be deciphered.

## Conclusion

This is the first study to elucidate the role of the non-iNKTs in atherosclerotic *Apoe^-/-^* mice and hyperlipidemic wild-type mice, opening the door for further scientific advancement in understanding the NKT cell biology and atherosclerosis progression. We observed that the absence of the *Apoe* gene in *Apoe^-/-^* mice significantly increases the number and activation of the non-iNKT cells. Our report highlights that the *Apoe* gene is crucial for the systemic immune modulation of the NKT subsets and warrants the study of NKT cells using the *Apoe^-/-^* mice, the gold standard model for atherosclerosis.

## Data Availability

All the data will be available and can be found on the journal website.

## Acknowledgments

We thank the NIH Tetramer Core Facility (contract number 75N93020D00005) for providing fluorophore-tagged PBS-57-loaded mouse CD1d tetramers.

## Funding

The work is partially supported by SERB Grant (CRG/2020/001641) by the Department of Science and Technology, Government of India.

## Disclosure

The authors declare that they have no known competing financial interests or personal relationships that could have appeared to influence the work reported in this paper.

## Author Contributions

R.C. – Conceived and designed research, performed experiments, analyzed data, interpreted results of experiments, prepared figures, drafted manuscript. S.D. - interpreted results of experiments. P.C.S. - Conceived and designed research, interpreted results of experiments, edited and revised manuscript, approved the final version of the manuscript.

## Notes

### Competing Interest Statement

The authors have declared no competing interest.

